# Efficient feature selection on gene expression data: Which algorithm to use?

**DOI:** 10.1101/431734

**Authors:** Michail Tsagris, Zacharias Papadovasilakis, Kleanthi Lakiotaki, Ioannis Tsamardinos

## Abstract

**Background:** Feature selection seeks to identify a minimal-size subset of features that is maximally predictive of the outcome of interest. It is particularly important for biomarker discovery from high-dimensional molecular data, where the features could correspond to gene expressions, Single Nucleotide Polymorphisms (SNPs), proteins concentrations, e.t.c. We evaluate, empirically, three state-of-the-art, feature selection algorithms, scalable to high-dimensional data: a novel generalized variant of OMP (gOMP), LASSO and FBED. All three greedily select the next feature to include; the first two employ the residuals resulting from the current selection, while the latter rebuilds a statistical model. The algorithms are compared in terms of predictive performance, number of selected features and computational efficiency, on gene expression data with either survival time (censored time-to-event) or disease status (case-control) as an outcome. This work attempts to answer a) whether gOMP is to be preferred over LASSO and b) whether residual-based algorithms, e.g. gOMP, are to be preferred over algorithms, such as FBED, that rely heavily on regression model fitting.

**Results:** gOMP is on par, or outperforms LASSO in all metrics, predictive performance, number of features selected and computational efficiency. Contrasting gOMP to FBED, both exhibit similar performance in terms of predictive performance and number of selected features. Overall, gOMP combines the benefits of both LASSO and FBED; it is computationally efficient and produces parsimonious models of high predictive performance.

**Conclusions:** The use of gOMP is suggested for variable selection with high-dimensional gene expression data, and the target variable need not be restricted to time-to-event or case control, as examined in this paper.

## 1 Background

The problem of feature selection (FS) has been studied for several decades in many data science fields, such as bioinformatics, statistics, machine learning, and signal processing. Given an outcome (or response) variable *γ* and a set *X* of *p* features (predictor or independent variables) of *n* samples, the task of FS is to identify the minimal set of features whose predictive capability on the outcome is optimal. A typical example of such a task is the identification of the genes whose expression allows the early diagnosis of a given disease.

Solving the FS problem has numerous advantages Tsamardinos and Aliferis (2003). Features can be expensive (or/and unnecessary) to measure, to store and process in the biological domains. For example, FS can reduce the cost of applying a diagnostic model by reducing the number of genes that will be measured. On top of that, parsimonious models are computationally cheaper and easier to visualize, inspect, understand and interpret. An FS algorithm of high quality often improves the predictive performance of the resulting model by removing the noise propagated by redundant features. This is especially true for models susceptible to the curse of dimensionality, perhaps one of the most common problems in biological datasets Lie (2014).

FS can also be used as a means of knowledge discovery and for gaining intuition on the data generation mechanisms. Indeed, there is theoretical connection between FS and the Bayesian (causal) network that describes best the data at hand Tsamardinos and Aliferis (2003). Following the Bayesian networks terminology, the Markov Blanket of a variable *γ* (time-to-event or disease status in this paper) is defined as the minimal set of variables that renders all other variables conditionally independent of *γ*. The Markov Blanket of *γ* carries all the necessary information about *γ*, and no other variable offers additional information about *γ* Niel et al. (2018). Under certain broad conditions the Markov Blanket has been shown to be the solution to the FS problem Tsamardinos et al. (2003). Identification of the Markov Blanket of the outcome variable is often the primary goal of FS and not the predictive model per se Borboudakis and Tsamardinos (2017). This is particularly true in bioinformatics, where, for example, the genes selected may direct future experiments and offer useful insight on the problem of interest, its characteristics and structure.

Over the years, there has been an accumulation of FS algorithms in many scientific fields. A recent review, along with open problems, regarding FS on high dimensional data, is provided in Saeys et al. (2007); Bühlmann and Van De Geer (2011); Boló-Canedo et al. (2014, 2016). The question that naturally arises is which algorithm to employ. Even though the answer is not known beforehand, the grounds upon which to decide, include computational efficiency, predictive performance, statistical soundness and correctness, and ability to handle numerous types of outcome variables. Based on these criteria we have selected three state-of-the-art algorithms, with desirable theoretical properties, scalable to high-dimensional data, to empirically evaluate them: a (novel) generalized variant of OMP (gOMP), LASSO and FBED. Tibshirani (1996) suggested, in the statistical literature, *Least Absolute Shrinkage and Selection Operation* (LASSO), while Pati et al. (1993); Davis et al. (1994) suggested, in the signal processing literature, *Orthogonal Matching Pursuit* (OMP). In this paper we propose the use of gOMP that generalizes to many types of outcome variables. *Forward-Backward with Early Dropping* (FBED) Borboudakis and Tsamardinos (2017) was recently introduced in the machine learning literature.

*In our empirical study, using real, publicly available gene expression data with time-to-event and case-control outcome variables, we demonstrate that gOMP is on par, or outperforms LASSO in several aspects.* gOMP produces predictive models of equal or higher accuracy, while selecting less features than LASSO. When comparing gOMP with FBED, similar conclusions were drawn. gOMP is highly efficient (computationally-wise), due to the fact that unlike FBED, which repeatedly fits regression models, it is residual-based and fits far less regression models. gOMP is on par with FBED in terms of predictive performance, selecting roughly unvarying number of features. In addition, gOMP and FBED’s method of selection is agnostic on the type of the outcome variable, rendering them more generalized FS methods. On the contrary, LASSO is heavily dependent on the outcome variable; each type of outcome variable requires different handling.

The rest of the paper is organized as follows; Section 2 briefly presents the three feature selection algorithms and discusses some of their properties. In Section 3 we describe the experimental setup and in Section 4 we present the results of the comparative evaluation of gOMP against LASSO and gOMP against FBED on gene expression data. A discussion on our results follows and the Conclusions alongside with future directions close the paper.

## 2 Methods

I this section we will briefly state the feature selection algorithms we will compare along with some key characteristics.

### 2.1 FBED

The oldest FS method is the forward regression (or forward selection) method; it repeatedly applies regression models in a greedy manner. At the first step, the outcome variable is regressed against every feature, selecting the feature producing the highest (statistically significant) association. In subsequent steps, the feature mostly (statistically significantly) associated with the outcome, given the already selected features, is selected. The process stops, when there is no statistically significant association between a feature and the outcome variable.

The forward regression lacks scalability to high-dimensional data. Forward-Backward with Early Dropping (FBED) Borboudakis and Tsamardinos (2017) is a recently proposed FS algorithm that overcomes this drawback. Compared to forward regression, the key element of FBED that makes it scalable to high dimensional data is that it removes, at every step, the non significant features (Early Dropping heuristic). FBED’s final step includes a backward selection to remove falsely selected features. Borboudakis and Tsamardinos (2017) showed that FBED identifies a subset of the outcome variable’s Markov Blanket, namely the parents and children in Bayesian network terminology. Iterating the algorithm one more time, by retaining the selected features and considering the features previously removed, FBED is able to identify the full Markov Blanket of the outcome variable. This holds true provided that the distribution of the data can be faithfully represented by a Bayesian network.

### 2.2 gOMP

Orthogonal Matching Pursuit (OMP) Chen et al. (1989); Pati et al. (1993); Davis et al. (1994) is a greedy forward-search algorithm. The procedure is exactly the same as in forward regression method. Their main difference is that OMP works with residuals and hence is highly efficient. At the initial step the feature (*x*_*j*_) having the largest absolute correlation with the outcome variable is selected and a regression model is fitted. The regression model produces a residual vector orthogonal to *x*_*j*_, now considered to be the outcome. New correlations between the updated outcome and the remaining features are calculated. The feature with the largest absolute correlation is then selected. At every step, the regression model with the candidate feature (including all previously selected features) is fitted and the norm of the residuals is computed. The process stops when the norm is below a pre-specified threshold.

In the established OMP algorithm, the norm-based stopping criterion cannot be defined beforehand conveniently. To overcome this, OMP runs multiple times. Our proposed modification of OMP, called generalized OMP (gOMP), uses a completely different stopping criterion. When a new feature is chosen for possible inclusion, the model including the candidate feature is fitted and its log-likelihood is calculated. If the increase in the log-likelihood values between the current model and the previous model (without the candidate feature) is above a certain threshold value, the feature is selected, otherwise the process stops. Such a threshold value is the 95% quantile of the *χ*^2^ distribution, forming a hypothesis test at the 5% significance level. The use of this stopping criterion, generalizes OMP’s functionality on accepting numerous types of outcome variables, including multi-class, survival, left censored, counts and proportions to name a few, thus being able to handle various regression models.

### 2.3 LASSO

Least Absolute Shrinkage and Selection Operation (LASSO) Tibshirani (1996) is perhaps the most popular FS algorithm, owing much of its success to its computational efficiency. Similarly to OMP, it is residual-based, with an extra degree of complexity. LASSO is formulated around the penalized minimization of an objective function (usually log-likelihood), which depends upon the type of the outcome variable. The regression coefficients live in a constrained space. Their magnitude depends upon the value of the penalty parameter, which forces them to shrink towards zero, hence feature selection is performed automatically.

### 2.4 A brief discussion of the gOMP, FBED and LASSO algorithms

Residual-based algorithms share a common statistical theoretical background. For a given statistical model, produced by regressing the outcome variable on a set of features, correlation among the resulting residual vector and a new feature, indicates correlation between that feature and the outcome variable. OMP and LASSO being residual-based algorithms, select the next best feature using residuals and fit substantially fewer regression models, rendering them computationally efficient. The complexity, or number of operations, required by LASSO is 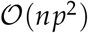 Rosset and Zhu (2007), where *n* and *p* denote the sample size and number of features, respectively. The number of calculations (regression models and correlations) gOMP performs is *p* (*s* + 1) where *s* denotes the number of selected features.

FBED approaches the task of FS from the Bayesian networks perspective. Forward selection fits *p* (*s* + 1) regression models where *p* is the number of variables and *s* the number of selected variables. On the other hand, backward selection fits *p* + (*p* − 1) + … + (*p* − *s*) regression models. FBED’s time complexity is bounded (worst case scenario) by the sum of these regression models. In practice though, its complexity is substantially lower, but cannot be quantified analytically, as this depends upon the input data Borboudakis and Tsamardinos (2017).

Blumensath and Davies (2007) described some differences between OMP and the standard forward selection method. The main difference mentioned in their work, which is in fact the same difference between OMP and FBED, is that OMP selects the new feature based on the inner product, i.e. the angle between the features and the current residual. On the contrary, FBED (and similar model fitting algorithms) will select the feature with the smallest angle, followed by the projection of this feature onto the orthogonal subspace in such a way so as to maximize an objective function, e.g. log-likelihood.

OMP is very similar to the residual-based *forward stepwise regression* Weisberg (1980); Efron et al. (2004). The difference is that in the latter, a feature is selected if its correlation with the current residual is statistically significant, or if its magnitude is above a certain threshold Stodden (2006).

In statistics, consistent model selection translates into selecting the correct features with probability tending to 1 as the sample size tends to infinity. Zhang (2009) discussed the necessary and sufficient conditions for a greedy least squares regression algorithm (such as gOMP) to select features consistently. These conditions match the necessary and sufficient conditions for LASSO mentioned by Zhao and Yu (2006). In practice though, LASSO tends to choose considerably more features than necessary, leading to an abundance of falsely selected features (false positives) and hence model consistency is not guaranteed. On the other hand, gOMP (and FBED) produces more parsimonious models. Borboudakis and Tsamardinos (2017) showed that under specific conditions, FBED identifies the minimal set of features with the maximal predictive performance (Markov Blanket).

## 3 Experiments on gene expression data

The first goal of this paper is to make suggestions as to which algorithm, gOMP or LASSO is more suitable for gene expression data analysis. By contrasting gOMP with FBED, the second goal is to provide some suggestions as to whether residual-based FS algorithms, or regression model based FS algorithms should be applied in practice. In order to compare each pair of algorithms on equal grounds, we implemented two distinct cross-validation procedures for each pair of comparisons, gOMP-LASSO and gOMP-FBED. The decision was based on the grounds of each algorithm having a different number of tunable hyper-parameters, as discussed in Section 3.4.

We used the packages *MXM* Lagani et al. (2017) for the gOMP and FBED algorithms, while for LASSO we used the R package *LASSO* Friedman et al. (2010). The R packages utilized for the predictive models are outcome-dependent and are mentioned below.

Hastie et al. (2017) performed a simulation study comparing LASSO with other competing algorithms by using simulated datasets only. They concluded that LASSO outperformed two state-of-the-art FS algorithms, forward stepwise regression Weisberg (1980) and best-subset-selection Bertsimas et al. (2016). On the contrary, Borboudakis and Tsamardinos (2017) compared FBED with LASSO using real high dimensional data from various fields, biology, text mining and medicine. Their results showed that LASSO produced predictive models with higher performance at the cost of being more complex. Borboudakis and Tsamardinos (2017) concluded that there is no clear winner between LASSO and FBED and the choice depends solely on the goal of the analysis.

The drawback of simulated data is that they do not necessarily portray the complex structure observed with real data. We have conducted extensive experiments focusing on real, publicly available, preprocessed gene expression data Lakiotaki et al. (2018).

### 3.1 The two outcome scenarios

We considered two different types of outcome variables: (censored [^1^ Censoring occurs when we have limited information about individual’s survival time, the exact time-to-event time is unknown.]) time-to-event and case-Control. For the time-to-event outcome scenario we investigated 6 gene expression datasets, most of which spanning to a few thousands of features (genes). For the case-control (binary) outcome scenario we investigated 12 gene expression datasets, most of which including more than 54, 000 features. Table 1 provides information about the datasets used in our experiments. The gene expression data (GSE annotated) are available for downloading from BioDataome Lakiotaki et al. (2018).

- **(Censored) time-to-event outcome**. The event of interest can be death, disease relapse, or any other time-related event. The goal of FS is to identify the subset of features, e.g. genes, mostly correlated with the survival time. All FS algorithms adopted a Cox proportional hazards model. LASSO using Cox proportional hazard regression model was suggested by Park and Hastie (2007). On the contrary, this is the first work where time-to-event outcome is treated by gOMP. Other survival regression models include the Weibull, log-logistic, exponential and log-normal model, for which, unlike LASSO, gOMP can easily be adapted to them. For gOMP, the martingale residuals Therneau et al. (1990) were calculated, while other options include the deviance, score, and Schoenfeld residuals.
- **Case-control outcome** [^2^ In this work, we focused on unmatched case-control gene expression data.], is frequently met in gene expression data. FS aims at identifying the minimal subset of genes that best discriminates among two classes. All three FS algorithms employed logistic regression. For gOMP in specific, similarly to Lozano et al. (2011), we calculated the raw residuals 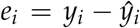, with other available options being the deviance and Pearson residuals.

**Table 1.** Information on the datasets used. The column *#Censored* refers to the number of samples whose value was right censored; there is limited information about individual’s survival time, the exact time-to-event time is unknown. The GSE annotation refers to the GEO accession number. All GSE annotated data are available for download from BioDataome.

| Time to event data |  |  |  |
| --- | --- | --- | --- |
| Dataset | #Samples | #Censored | #Features |
| Vijver Van De Vijver et al. (2002) | 295 | 207 | 70 |
| Veer Van't Veer et al. (2002) | 78 | 44 | 4,751 |
| Bullinger Bullinger et al. (2004) | 116 | 49 | 6,283 |
| Beer Beer et al. (2002) | 86 | 62 | 7,129 |
| Ros.2002 Rosenwald et al. (2002) | 240 | 102 | 7,399 |
| Ros.2003 Rosenwald et al. (2003) | 92 | 28 | 8,810 |

Table 1: Information on the datasets used.
| Binary outcome data |  |  |  |
| --- | --- | --- | --- |
| Dataset | #Samples | Class distribution (%) | #Features |
| Breast cancer Wang et al. (2005) | 286 | (99%, 1%) | 17,816 |
| GSE8006 | 206 | (89%, 11%) | 54,675 |
| GSE58331 | 175 | (83%, 17%) | 54,675 |
| GSE53987 | 205 | (73%, 27%) | 54,675 |
| GSE52724 | 286 | (49%, 51%) | 54,675 |
| GSE50705 | 351 | (84%, 16%) | 54,675 |
| GSE40791 | 194 | (48%, 52%) | 54,675 |
| GSE32474 | 174 | (85%, 15%) | 54,675 |
| GSE31210 | 246 | (92%, 8%) | 54,675 |
| GSE21545 | 223 | (57%, 43%) | 54,675 |
| GSE16214 | 240 | (66%, 34%) | 54,675 |
| GSE5281 | 161 | (54%, 46%) | 54,675 |

### 3.2 Evaluation pipeline

We employed a fully-automated machine learning pipeline for assessing the performance of each FS method. For this, an analysis pipeline similar to Just Add Data Bio or JAD Bio, trademark of Gnosis Data Analysis (gnosisda), was applied, ensuring methodological correctness.

We conducted an 8-fold cross-validation for time-to-event outcome and a 10-fold cross-validation for case-control outcome data. In the time-to-event outcome scenario the data were randomly allocated in the 8 folds. For the case-control outcome scenario stratified random sampling ensured that the ratio of the number of cases to the number of controls was kept nearly the same within each of the 10 folds. In each fold, the three algorithms were tested using a range of hyper-parameters (depending on the algorithm). Advanced linear and non linear statistical models and machine learning algorithms were employed in order to assess the predictive performance of the features selected by gOMP, LASSO and FBED.

### 3.3 Bias correction of the estimated performance

The estimated predictive performance of the best model is optimistically biased (the performance of the chosen model is overestimated) and a bootstrap-based bias correction method was applied Tsamardinos et al. (2018).

Upon completion of the cross-validation, the predicted values produced by all predictive models across all folds is collected in a matrix *P* of dimensions *n* × *M*, where *n* is the number of samples and *M* the number of trained models or configurations. Sampled with replacement a fraction of rows (predictions) from *P* are denoted as the in-sample values. On average, the newly created set will be comprised by 63.2% of the original individuals [^3^ The probability of sampling, with replacement, a sample of *n* numbers from a set of *n* numbers is 1 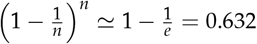], whereas the rest 36.8% will be random copies of them Efron and Tibshirani (1994). The non re-sampled rows are denoted as out-of-sample values. The performance of each model in the in-sample rows is calculated and the model (or configuration) with the optimal performance is selected, followed by the calculation of performance in the out-of-sample values. This process is repeated *B* times and the average performance is returned.

Note, that the only computational overhead is with the repetitive re-sampling and calculation of the predictive performance, i.e. no model is fitted nor trained. The final estimated performance usually underestimates the true performance, but this negative bias is smaller than the optimistic uncorrected performance.

### 3.4 Tuning the feature selection algorithms

When comparing gOMP to LASSO, 10 threshold values for gOMP were chosen, spanning from 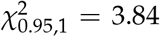 (the 95% upper quantile of the *χ*^2^ distribution with 1 degree of freedom), up to 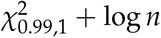 . Alternatively, this represents a range of different critical values corresponding to different significance levels, e.g. 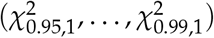. The existence of continuous features only justifies a single degree of freedom. The term log *n* implies that BIC Schwarz et al. (1978) is applied instead. As for LASSO, the default number of values of *λ* is 100, starting from 0 up to a data dependent maximum value. For a fair comparison with gOMP, we selected 10 values equally spaced within that range.

FBED has fewer tunable hyper-parameter values. The selection process in FBED can be based upon the extended BIC (eBIC) Chen and Chen (2008), or the Likelihood ratio test. BIC can be used as a special case of the eBIC, while for the likelihood ratio test the significance level was chosen to be 0.01 and 0.05. As for the number of runs of FBED, Borboudakis and Tsamardinos (2017) showed that a single re-run of the algorithm is usually sufficient. In order to match the number of FBED’s hyper-parameter values, we selected 4 threshold values for gOMP, 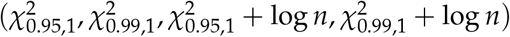. The first two correspond to log-likelihood ratio test values at 0.01 and 0.05 significance level, whereas the remaining two correspond to using BIC.

### 3.5 Predictive algorithms

- **Support Vector Machines (SVM)** were proposed by Cortes and Vapnik (1995). The key idea is to transfer the data into higher dimensions by using kernel functions. SVM for survival outcome variables with censored values first appeared in Shivaswamy et al. (2007) and were based on a simple modification upon the SVM’s constraint-optimization formulation. We applied a Gaussian kernel (also called Radial Basis Function) and selected *9* values for its tuning parameter. SVM are available in the the R packages *survivalsvm* Fouodo (2018) and *e1071* Meyer et al. (2017) for the time-to-event outcome and the case-control outcome, respectively.
- **Random Forests (RF)** were proposed by Breiman (2001). RF perform Decision Trees Breiman et al. (1984) repeatedly and aggregate their results. Random Forests for time-to-event outcomes were proposed later by Ishwaran et al. (2008). RF require the specification of the parameter *nsplit*, i.e. the number of splitting points to be evaluated for each feature; higher values of *nsplit* may lead to more accurate predictions at the expense of greater computational load. We selected *3* values while setting the remaining parameters to their default values. We used the R package *randomForestSRC* Ishwaran and Kogalur (2017) which can handle both case-control (binary) and time-to-event (survival) outcome variables.
- **Ridge regression (RR)** Hoerl and Kennard (1970) shrinks the regression coefficients by imposing a penalty on the sum of their squared values [^4^ LASSO imposes the penalty on the sum of their absolute values, (see Eq. (1) in the Supplementary material.]. The difference with LASSO is that the regression coefficients do not shrink towards zero. For both time-to-event and case-control outcomes we selected a set of *11* values for the shrinkage parameter. We used the R packages *survival* Therneau (2017) and *glmnet* Friedman et al. (2010) for the time-to-event outcome and the case-control outcome respectively.

For the gOMP-LASSO comparison, 230 models were trained for each scenario. For each of the 10 hyper-parameter values of gOMP and LASSO, 23 configurations, 9 SVM configurations, 3 RF configurations and 11 RR configurations were tested. For the gOMP-FBED comparison 92 predictive models were produced (23 configurations for each of the 4 hyper-parameters of gOMP and FBED).

### 3.6 Predictive performance metrics and number of selected features

For time-to-event outcome, the concordance index (C-index) Harrell et al. (1996) is the standard performance metric for model assessment in survival analysis. It expresses the probability that, for a pair of randomly chosen samples, the sample with the highest risk prediction will be the first one to experience the event (e.g death). C-index measures the percentage of pairs of subjects correctly ordered by the model in terms of their expected survival time. A model ordering pairs at random (without use of any feature) is expected to have a C-index of 0.5, while perfect ranking would lead to a C-index of 1. When there are no censored values, the C-index is equivalent to the Area Under the Curve (AUC).

AUC was employed as the performance metric in the case-control outcome scenario. AUC represents the probability of correctly classifying a sample to the class it belongs to, thus takes values between 0 and 1, where 0.5 denotes random assignment. Unlike the accuracy metric (proportion of correctly classified samples), AUC is not affected by the distribution of the two classes (cases and controls).

We also examined the number of features each algorithm selected. As mentioned in the Introduction, the Markov Blanket of the outcome variable is the minimal subset of features, and often the primary goal of FS is to identify these features per se.

We statistically evaluated the differences in predictive performance and number of selected features. Due to the small number of datasets used in our empirical study, we calculated the p-value using permutations Pesarin (2001). The differences in predictive performance (and number of selected features) between gOMP & LASSO and between gOMP & FBED are calculated and their average difference is used as the observed test statistic. Then, we randomly permuted, 1999 times, the sign of the differences, and each time calculated the mean difference. The p-value was computed as the proportion of times the permuted test statistics exceeded the value of the observed test statistic. This process was repeated 1000 in order to provide confidence intervals for the true p-value and thus provide safer conclusions. The same exact procedure was repeated, but instead of the differences, we used the logarithm of the ratios of the performances and the logarithm of the number of selected features.

### 3.7 Computational efficiency of the algorithms

The computational cost of each algorithm was measured during the 8 or 10-fold cross-validation. gOMP selects most features when its stopping criterion is low; 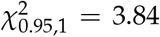 in our case. On the other hand, LASSO was applied with 10 *λ* values. FBED’s computational requirements were measured only when log-likelihood ratio test was applied with significance level equal to 0.05. All experiments were performed on an Intel Core i5-4690K CPU@3.50GHz, 32GB RAM desktop and the computational time was measured in seconds.

## 4 Results

Overall, as depicted in Figure 3, feature selection constitutes an integral part in gene expression data analysis for biomarker discovery. All three FS algorithms exhibit high predictive performance, however, the linear combination of features selected by gOMP and FBED is able to more clearly separate the data on the first two principal component space.

### 4.1 gOMP-LASSO: predictive performance and number of selected features

Table 2 summarizes the results of the comparison between gOMP and LASSO, while Figure 1 visualizes this information.

- **Time-to-event outcome scenario**. Figure 1(a) shows the relationship between the predictive performance (C-index) and the number of selected features. It is evident that in terms of predictive performance and number of selected features, gOMP outperforms LASSO, as it achieves better performance with a smaller set of features. The p-value for the mean difference in the C-index was equal to 0.045 with a 95% confidence interval (calculated using 1000 repetitions) equal to (0.0389, 0.058). The confidence interval shows that gOMP performed statistically significantly better than LASSO at a significance level of 6%. The results were similar for the mean log-ratio; the p-value was equal to 0.047, and its 95% confidence interval was (0.039, 0.057). LASSO selected st tistically significantly more features. The p-values for the mean difference and the mean log-ratio were less than 0.001 and so were their 95% confidence intervals.
- **Case-control outcome scenario**. The results for the binary outcome scenario are presented in Figure 1(b). LASSO achieved similar or better predictive performance than gOMP in most cases. LASSO though selected significantly more features than gOMP. In the worst case scenario, LASSO selected 10 times more features than gOMP. The p-value for the mean difference in the AUC was equal to 0.987, with a 95% confidence interval equal to (0.982, 0.992). The p-value for the mean log-ratio was equal to 0.989 with a 95% confidence interval equal to (0.985, 0.993). Both p-values and their corresponding confidence intervals indicate that the differences in the predictive performance of gOMP and LASSO, based on all 12 datasets are not statistically significant. LASSO selected statistically significantly more features than gOMP. Both p-values (referring to the mean difference and mean log-ratio) and their corresponding confidence intervals were less than 0.001%.

**Table 2:**
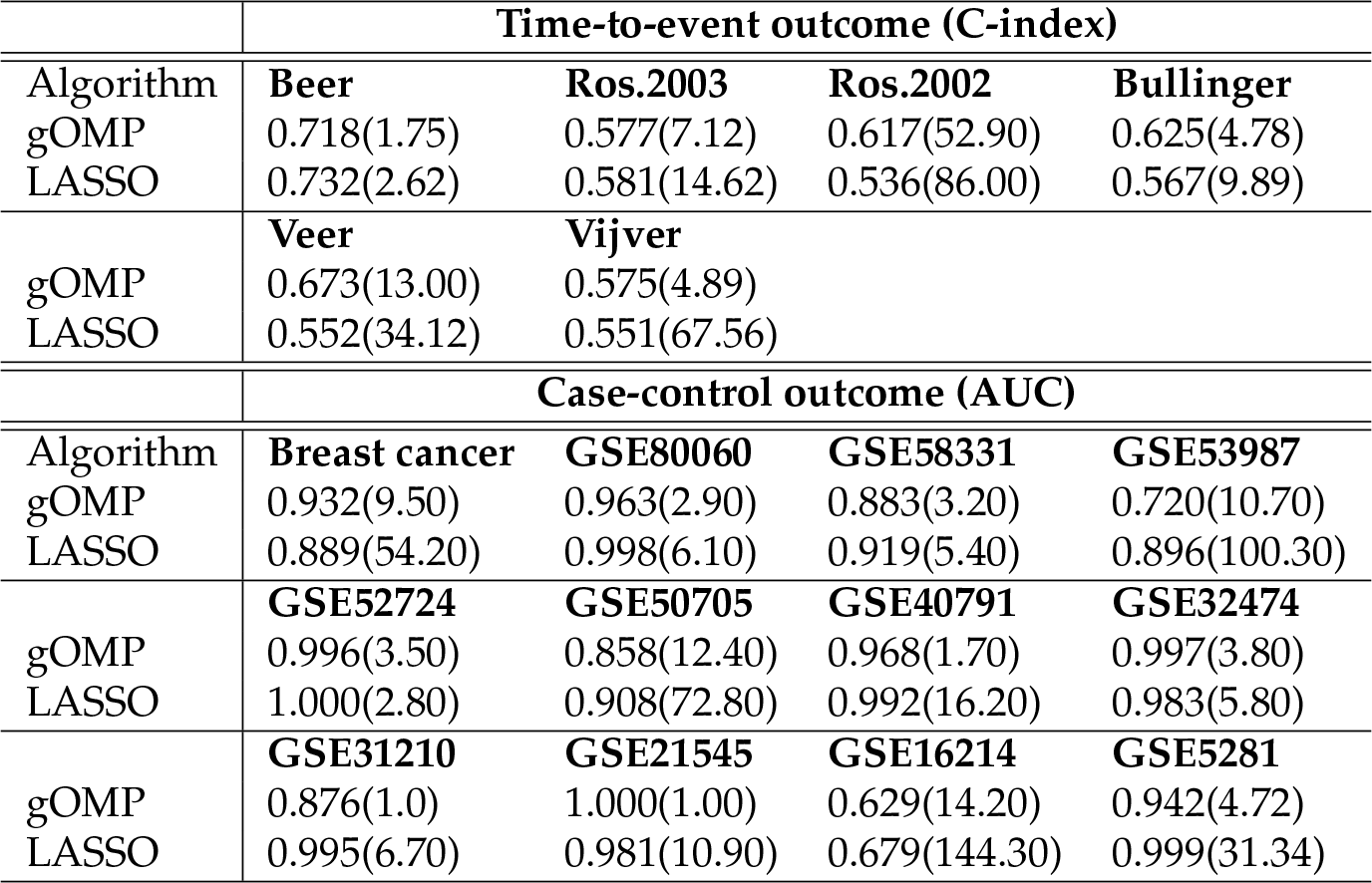
gOMP-LASSO: Predictive performance of gOMP and LASSO. The C-index was used in the time-to-event outcome scenario and the AUC in the case-control outcome scenario. Both metrics take values between 0 and 1, with high values being desirable. The average number of selected features appears inside the parenthesis.

**Table 3.**
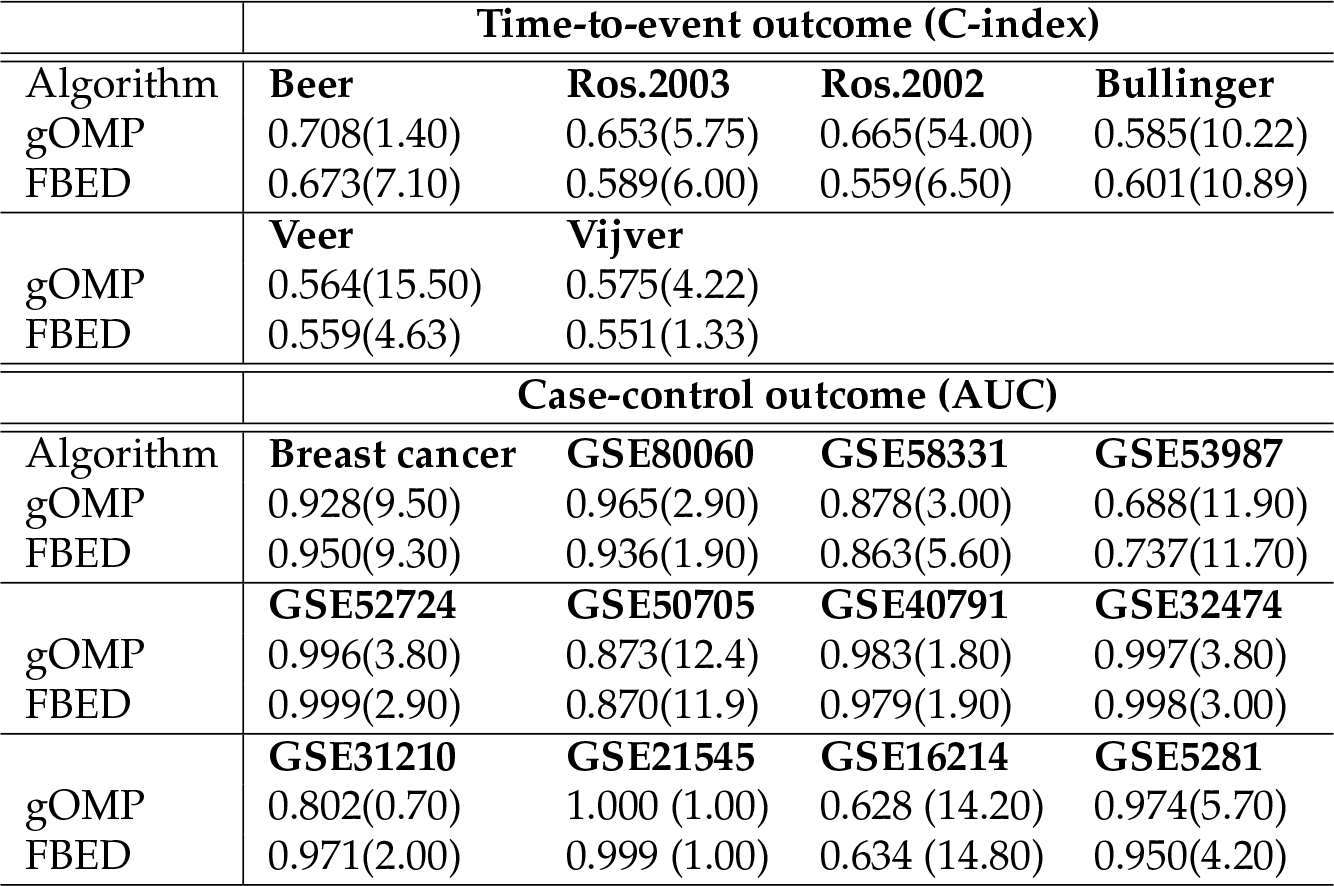
gOMP-FBED: Predictive performance of gOMP and FBED. The C-index was used in the time-to-event outcome scenario and the AUC in the case-control outcome scenario. Both metrics take values between 0 and 1, with high values being desirable. The average number of selected features appears inside the parentheses.

### 4.2 gOMP-FBED: predictive performance and number of selected features

The results of the comparison between gOMP and FBED are presented in Table 3 and Figure 2.

- **Time-to-event outcome scenario**. Figure 2(a) shows the relationship between the predictive performance (C-index) and the number of selected features. In terms of predictive performance, the results appear to be similar. The permutation based p-values for both the mean difference and the mean log-ratio of the predictive performance resulted in p-values equal to 0.032 with nearly the same 95% confidence intervals, equal to (0.024, 0.04). gOMP was selecting statistically significantly more features at the 7% significance level (p-value equal to 0.058), with a 95% confidence interval equal to (0.048, 0.069). gOMP was selecting, on average, 54 features in the dataset *Ros.2002*. Excluding this dataset, the 95% confidence intervals were shifted towards higher values: (0.053, 0.074) for the mean difference and mean log-ratio of the predictive performance, and (0.214, 0.250) for the number of selected features.
- **Case-control outcome scenario**. The results are presented in Figure 2(b). The permutation based p-values for the predictive performance show that there is no statistically significant difference. The p-value for the mean AUC difference was equal to 0.769, with a 95% confidence interval equal to (0.750, 0.790). The p-value for the mean AUC log-ratio was equal to 0.794 with a 95% confidence interval equal to (0.777, 0.812). Both p-values and their corresponding confidence intervals show that the predictive performance of gOMP and FBED, based on all 12 datasets are not statistically significantly different. The permutation based p-values for the difference in the number of selected features again showed no statistically significant difference. The p-value for the mean difference was equal to 0.497, with a 95% confidence interval equal to (0.475, 0.520). The p-value for the log-ratio was equal to 0.369 and its associated 95% confidence interval equal to (0.348, 390).

**Table 4.**
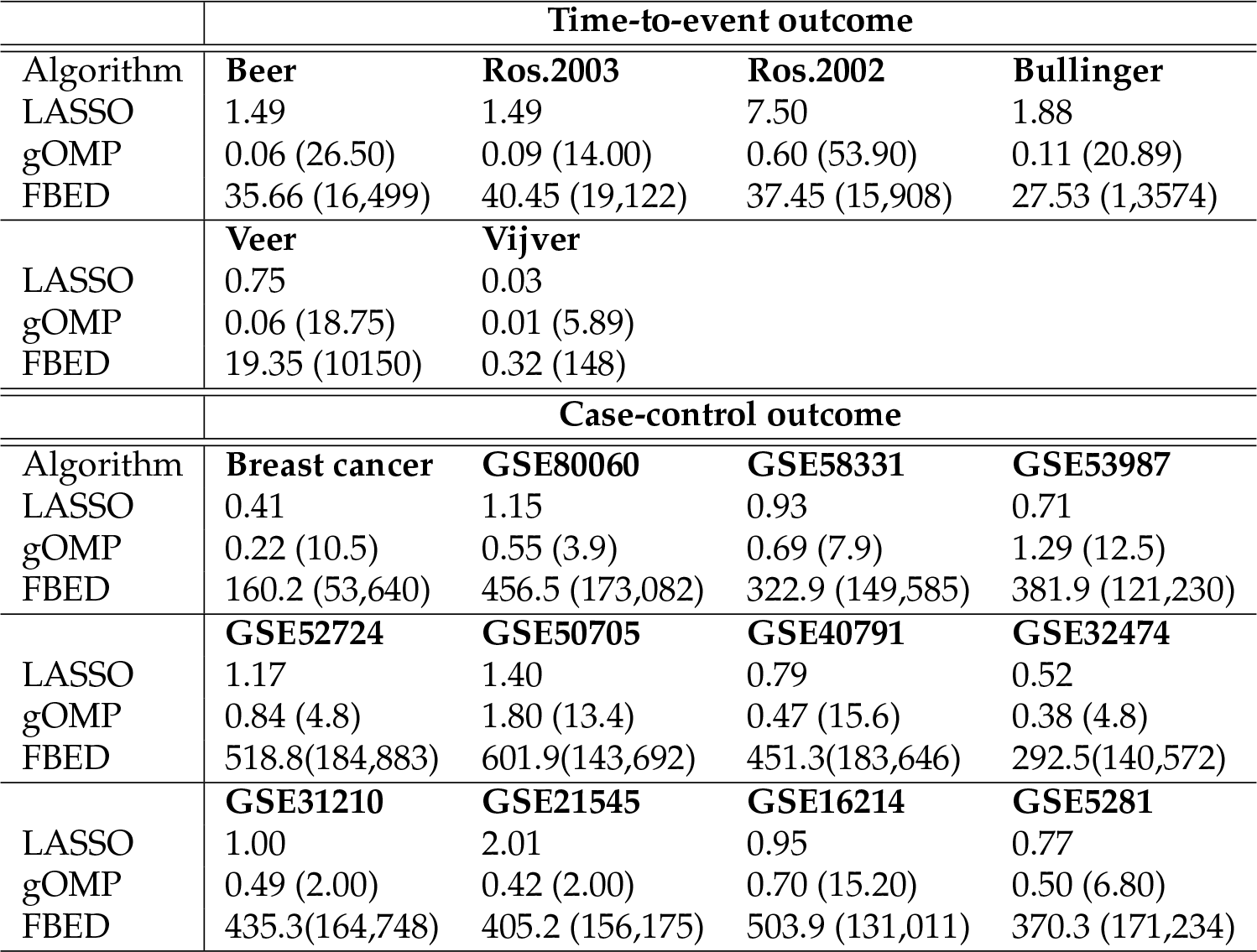
Average computational time (aggregated over all folds), in seconds, for each algorithm. The average number of regression models each algorithm fitted appears inside the parenthesis. For LASSO, the number of regression models fitted is not available, as this number depends upon the number of *λ* (hyper-parameter) values used, and hence is data independent.

**Table 5.** Average number of associated diseases for the gene sets selected by each algorithm. The number of selected genes is given in parenthesis. The lower number of associated diseases is italicized.

|  | Case-control outcome |  |  |  |
| --- | --- | --- | --- | --- |
| Algorithm | GSE80060 | GSE58331 | GSE53987 | GSE52724 |
| LASSO | 25.0 (5) | 26.0 (5) | 39.8 (121) | 92.0 (3) |
| gOMP | 5.5 (3) | 1.0 (3) | 77.7 (9) | 14.0 (4) |
| FBED | 49.0 (2) | 45.6 (8) | 28.4 (12) | 15.0 (3) |
|  | GSE50705 | GSE40791 | GSE32474 | GSE31210 |
| LASSO | 45.2 (78) | 69.1 (13) | 259.3 (6) | 12.8 (7) |
| gOMP | 8.2 (14) | 102.0 (2) | 596.5 (4) | - (1) |
| FBED | 6.3 (12) | 217.0 (2) | 444.5 (3) | 14.0 (2) |
|  | GSE21545 | GSE16214 | GSE5281 |  |
| LASSO | 101.0 (12) | 39.1 (158) | 67.2 (38) |  |
| gOMP | 77.0 (1) | 39.1 (16) | 33.7 (4) |  |
| FBED | 2.0 (1) | 56.0 (16) | 11.7 (4) |  |

### 4.3 Computational efficiency of the algorithms

gOMP and FBED are implemented in R, whereas LASSO is implemented in Fortran. Due to the nature of the algorithm, FBED fits substantially more regression models than gOMP. In addition, the standardization of the data, reduces the calculation of the correlation coefficients to calculation of inner products. On the other hand, fitting a logistic regression model involves repeated matrix multiplications.

For the time-to-event outcome scenario, gOMP is, on average, 9 times faster than LASSO, whereas for the case-control outcome scenario, gOMP is only 1.3 times faster. FBED is, on average, 162 times slower than gOMP and 18 times slower than LASSO in the time-to-event outcome scenario. In the case-control outcome scenario, FBED is 340 times slower than gOMP and 445 times slower than LASSO. Detailed information on the time comparisons is given in Table 4.

### 4.4 From feature selection to biological interpretation

All case-control gene expression datasets in this work have been measured by Affymetrix Human Genome U133 Plus 2.0 Array (GPL570). In such expression arrays a probe is a short DNA sequence targeting a short region of a transcript. Probes are grouped into probesets which are designed to target the same transcript with multiple measurements. We applied all three FS algorithms on the probeset level and mapped the selected probes to gene symbols using both Ensembl and annotation files for GPL570 from Gene Expression Omnibus-GEO and keeping the most frequent gene symbol. To gain some biological sense of the selected features (genes) we downloaded all gene-disease associations from DisGenet, which integrates data from expert curated repositories, GWAS catalogs, animal models and the scientific literature. The rationale behind this is that since gOMP and FBED almost always select a smaller number of predictive genes, we expect these genes to be more specialized genes for the outcome disease and thus be related to a smaller number of diseases compared to genes selected by Lasso. This is easily apparent in Table 5.

## 5 Discussion

We compared three feature selection algorithms, gOMP, LASSO and FBED, in survival regression and classification settings, using real gene expression data. An advantage of gOMP and FBED is that they can handle most kinds of regression models and types of outcome variables, whereas LASSO is not that flexible, i.e. each regression model requires the appropriate mathematical formulation. LASSO, on the other hand, due to its computational efficiency and predictive performance has gained research attention in various data science fields and numerous extensions and generalizations have been proposed over the years.

In this paper we showed that a reasonable alternative to LASSO is gOMP. Borboudakis and Tsamardinos (2017) showed that FBED achieves similar predictive performance to LASSO, at the cost of being computationally more expensive. We corroborated their results when comparing gOMP with FBED. We used gene expression data for predicting time-to-event and case-control outcome variables. To assess the quality of each FS algorithm we focused on three key elements, a) predictive performance, b) number of selected features and c) computational efficiency.

- **Predictive performance and number of selected features:** OMP leads to predictive models with higher predictive accuracy than LASSO in the time-to-event outcome scenario. With case-control outcome variables, LASSO and gOMP performed statistically equally well. LASSO, tends to select more features than necessary, leading to more complex predictive models. This has two disadvantages: a) the models are more difficult (and computationally expensive) to tune, train and interpret, b) if the primary goal of the re-searcher, or practitioner is to identify the relevant features (identification of the Markov Blanket per se), then gOMP should be preferred, as LASSO will have selected extra, re-dundant features. gOMP is on par with FBED in terms of predictive performance and number of selected features. Both of them select, on average, a similar number of features, produce parsimonious models and have similar predictive performance.
- **Computational efficiency:** gOMP and LASSO, being residual-based algorithms, are significantly more efficient than algorithms such as FBED, which fit substantially fewer regression models. In the time-to-event outcome scenario, gOMP was 9 times faster than LASSO, while in the case-control outcome scenario gOMP was only 30% faster than LASSO. We highlight also that computational cost is language or software dependent. Borboudakis and Tsamardinos (2017) showed that when using Matlab, the computational cost of FBED compared to LASSO is not always significant. gOMP and FBED accepting many diverse types of outcomes have been implemented in the R package *MXM* Lagani et al. (2017).

**Figure 1:**
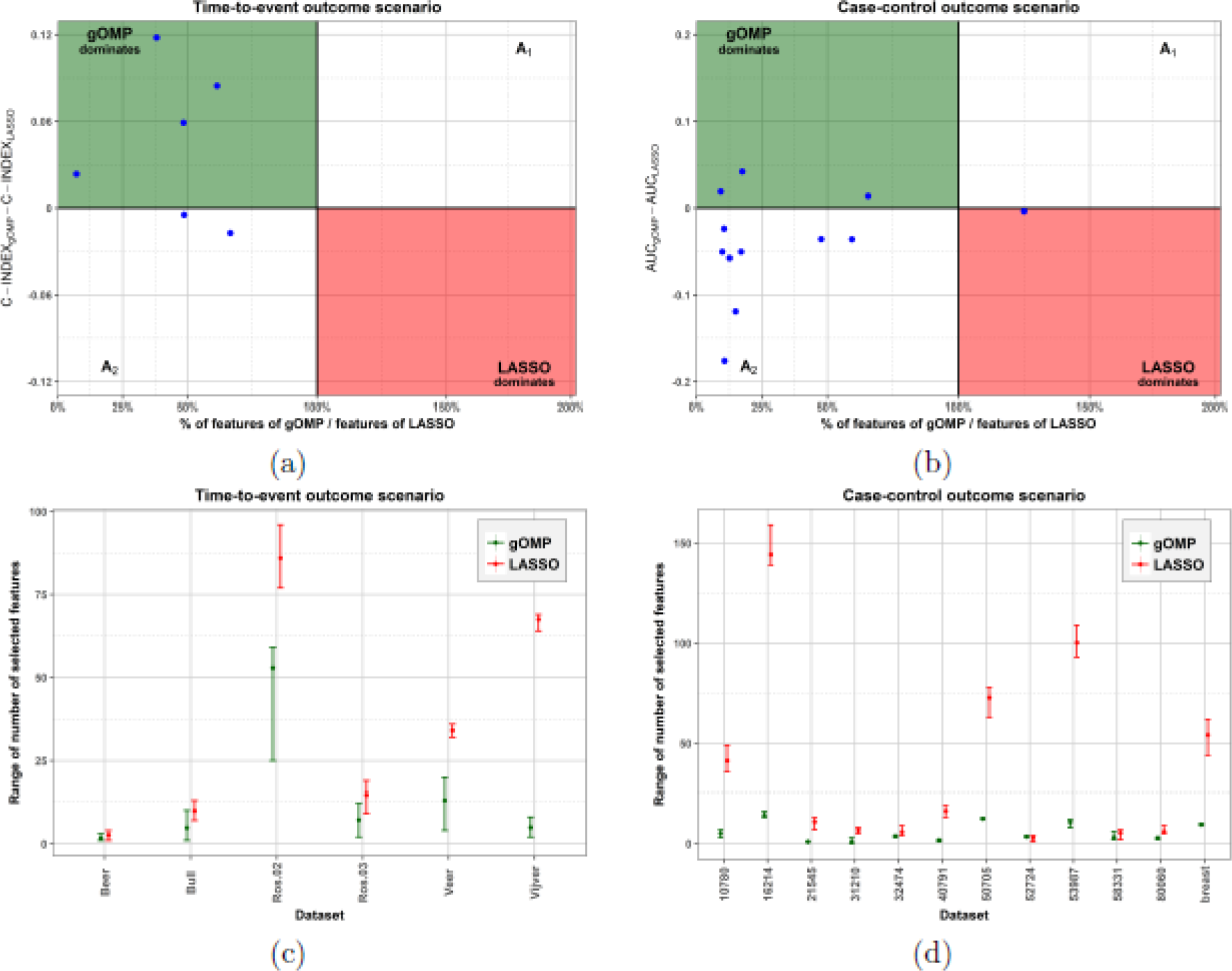
gOMP-LASSO. (a) & (b): Predictive performance vs number of selected features. The *x*-axis represents the percentage-wise ratio of selected features between gOMP and LASSO. Values less than 100% indicate that gOMP selected fewer features than LASSO. The *y*-axis represents the predictive performance difference between gOMP and LASSO, with positive values indicating that gOMP performs better than LASSO. In the top left quartile, shown in green, gOMP outperforms LASSO in both performance and number of selected features, whereas LASSO outperforms gOMP in both performance metrics in the bottom right quartile (red area). *A*_1_: gOMP has better predictive performance than LASSO while selecting more features. *A*_2_: LASSO has better predictive performance than gOMP while selecting more features.(c) & (d): Range of the number of selected features for each dataset. The solid circle is the average, and the vertical lines denote the minimum and maximum number of the selected features over all folds.

**Figure 2:**
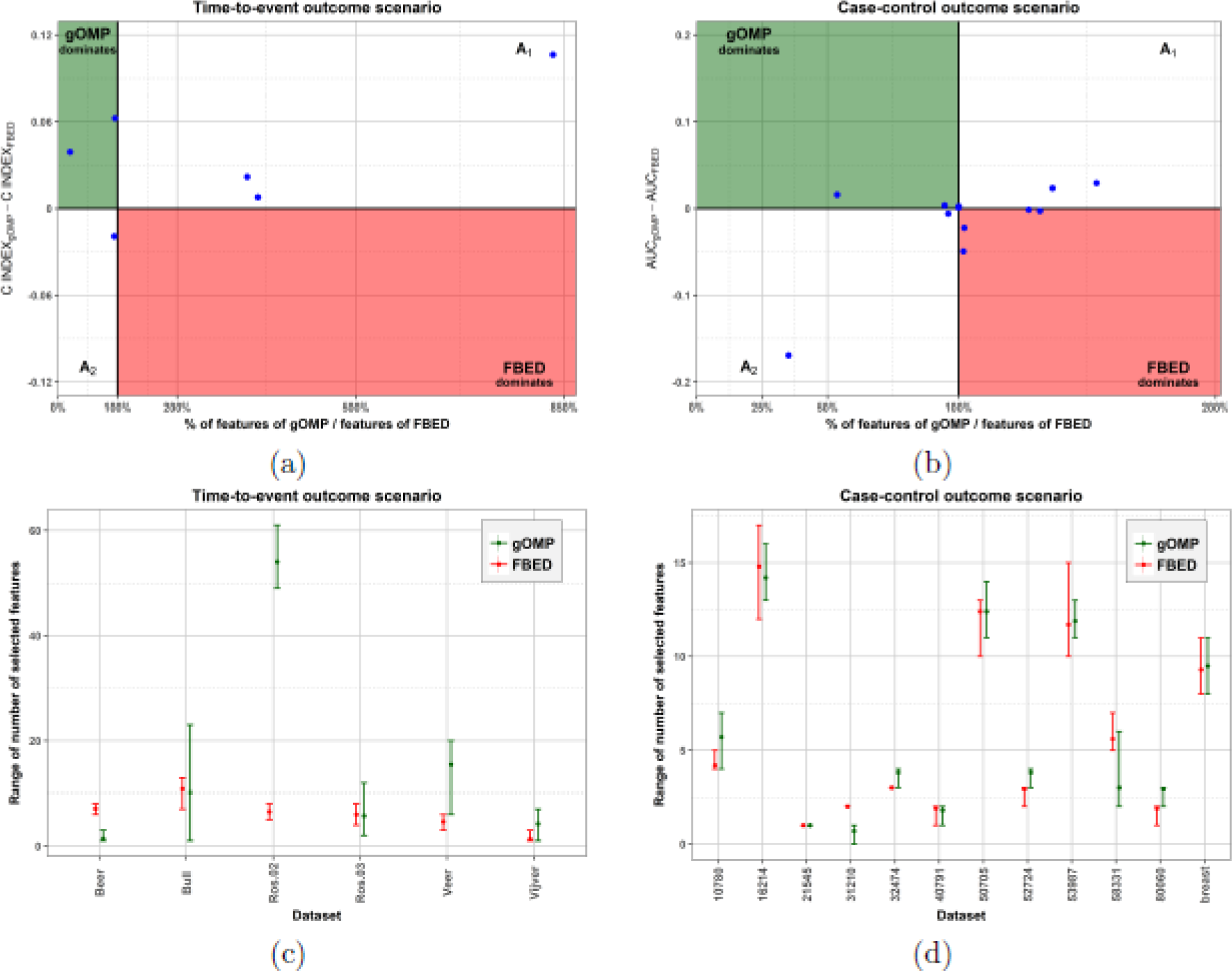
gOMP-FBED. (a) & (b): Predictive performance vs number of selected features. The *x*-axis represents the percentage-wise ratio of selected features between FBED and gOMP. Values less than 100% indicate that gOMP selected fewer features than FBED. The *y*-axis represents the difference in the predictive performance between gOMP and FBED, with positive values indicating that gOMP performs better than FBED. In the top left quartile, shown in green, gOMP outperforms FBED in both metrics, whereas FBED outperforms gOMP in both metrics in the bottom right quartile (red area). *A*_1_: gOMP has better predictive performance than FBED while selecting more features. *A*_2_: FBED has better predictive performance than gOMP while selecting more features. (c) & (d): Range of the number of selected features for each dataset. The solid circle is the average, and the vertical lines denote the minimum and maximum number of the selected features over all folds.

**Figure 3:**
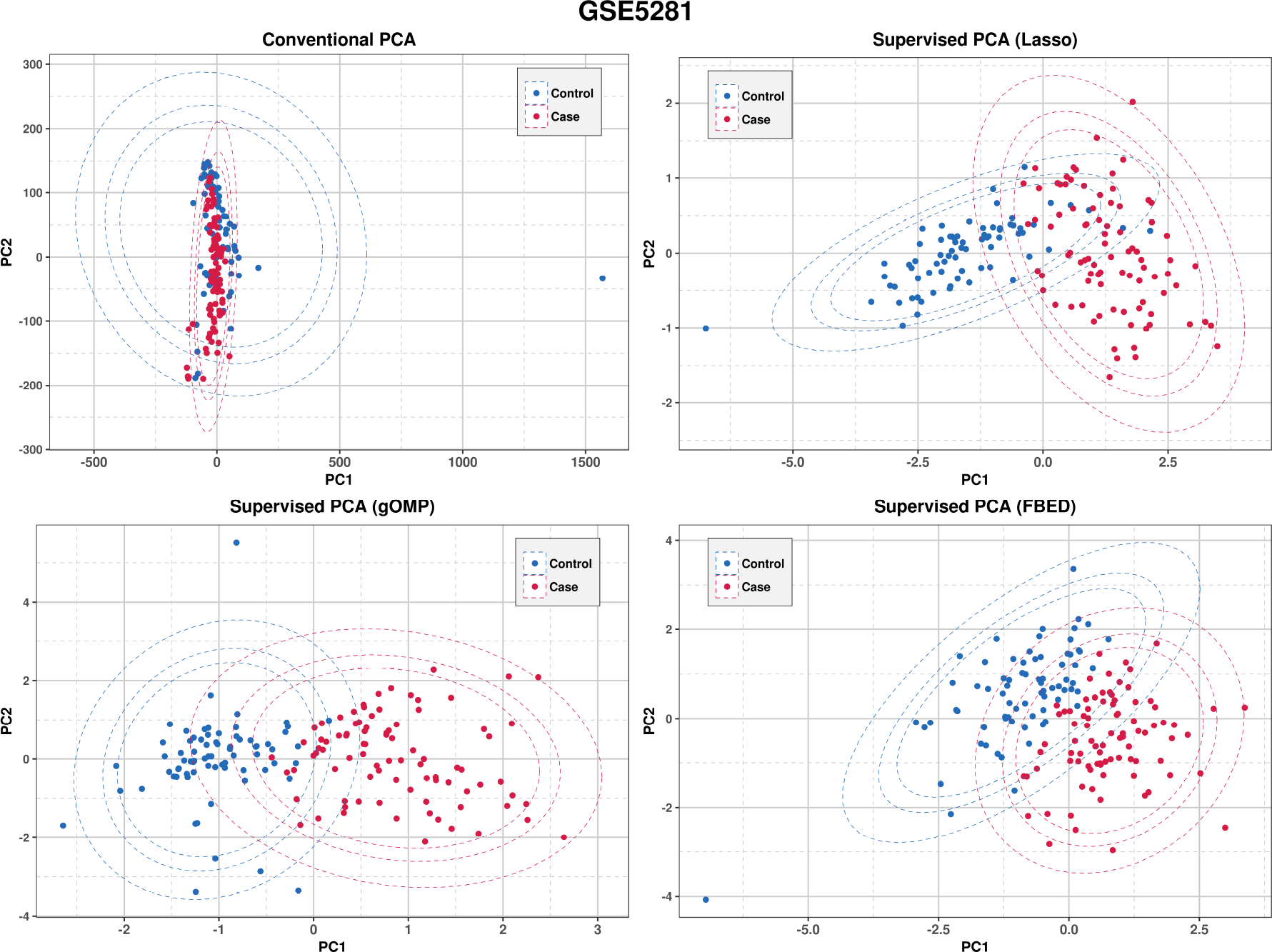
The first two principal components using all features (conventional PCA) and using the features selected by each algorithm (supervised PCA).

## 6 Conclusions

Our empirical study clearly points out that LASSO should not be the only FS algorithm to consider. LASSO is computationally efficient, with high predictive performance at the price of producing complex models by including many non relevant features. FBED is computationally more expensive, achieves high predictive performance and produces parsimonious predictive models. gOMP combines the benefits of both types of algorithms. Similarly to LASSO, gOMP tackles the FS problem from a geometrical standpoint (residual-based algorithm), thus, unlike FBED, it fits a smaller number of regression models. However, like FBED, gOMP produces parsimonious predictive models of high performance.

Ein-Dor et al. (2004) demonstrated that multiple, equivalent prognostic signatures for breast cancer can be extracted just by analyzing the same dataset with a different partition in training and test set, showing the existence of several genes which are practically interchangeable in terms of predictive power. We provided evidence of this phenomenon since gOMP, LASSO and FBED achieved, many times, similar performances by selecting different sets of features. SES Tsamardinos et al. (2012); Lagani et al. (2017) is one of the few FS algorithms that discovers statistically equivalent feature sets. More recently, Pantazis et al. (2017) proposed a solution for discovering equivalent sets of features using LASSO followed by computational geometry techniques. Ongoing research focuses on the identification of multiple sets of features that are statistically equivalent in terms of predictive performance, for both gOMP and FBED.

## Declarations

### Ethics and consent to participate

Not applicable.

### Consent for publication

Not applicable.

### Author’s contributions

MT has participated in the design of the study, wrote the code, performed the experiments and drafted the manuscript. ZP has contributed to the manuscript and to the design of the study. KL has contributed to the manuscript. IT contributed to the manuscript, and to the design of the study. All authors have read and approved the final manuscript.

### Availability of data and materials

All data are publicly available from the Gene Expression Omnibus database (GEO, http://www.ncbi.nlm.nih.gov/). The gene expression data (GSE annotated) are available for downloading from BioDataome. The gOMP and FBED algorithms, including more types of outcomes, are available in the R package MXM https://cran.r-project.org/web/packages/MXM/index.html.

## Acknowledgements

We would like to acknowledge Klio Maria Verrou for providing constructive feedback and Spyros Kotomatas for commenting on an earlier version of this paper.

## List of abbreviations

Not applicable.

## Authors Information Statement

MT

ZP

KL: IT

